# Syntaxin 17 promotes lipid droplet formation by regulating the distribution of acyl-CoA synthetase 3

**DOI:** 10.1101/124065

**Authors:** Hana Kimura, Kohei Arasaki, Yuki Ohsaki, Toyoshi Fujimoto, Mitsuo Tagaya

## Abstract

Lipid droplets (LDs) are ubiquitous organelles that contain neutral lipids and are surrounded by a phospholipid monolayer. How proteins specifically localize to the phospholipid monolayer of the LD surface has been a matter of extensive investigations. Here we show that syntaxin 17 participates in LD biogenesis by regulating the distribution of acyl-CoA synthetase 3 (ACSL3), a key enzyme for LD biogenesis that redistributes from the endoplasmic reticulum to LDs during LD formation. Time course experiments revealed that syntaxin 17 binds to ACSL3 in the initial stage of LD formation, and that ACSL3 is released as a consequence of competitive binding of SNAP23 to syntaxin 17 in the maturation stage. We propose a model in which ACSL3 redistributes from the endoplasmic reticulum to LDs through association with syntaxin 17 and SNAP23-mediated dissociation from syntaxin 17. We also provide evidence that lipid raft-like structures are important for LD formation and SNAREs-ACSL3 interactions.

## INTRODUCTION

Lipid droplets (LD) are ubiquitous organelles that store neutral lipids such as triacylglycerol (TAG) and sterol esters, and play central roles in energy and lipid metabolism (*Walther and Farese, 2012*). LDs are dynamic and diverse organelles, their size and number depending on cellular energy and the metabolic state, and their protein and lipid compositions varying with the cell type and the degree of LD maturation in individual cell types (*Ohsaki et al., 2014; Pol et al., 2014; Thiam and Beller, 2017*). Moreover, LDs are in contact with many organelles, including the endoplasmic reticulum (ER), mitochondria, and peroxisomes (*Gao and Goodman, 2015; Ohsaki et al., 2017*). Recent studies revealed that LDs have additional functions, such as in ER stress responses, protein storage, protein degradation, and viral replication (*Stordeur et al., 2014; Welte, 2015*).

LDs are unique among cellular organelles in that they are surrounded by a phospholipid monolayer. LD formation starts in the endoplasmic reticulum (ER) at pre-defined or random sites (*Kassan et al., 2013; Thiam and Forêt, 2016*), with the formation of lipid lenses of around 50 nm in the intermembrane space of the ER lipid bilayer (*Choudhary et al., 2015*). The formation of lenses and their enlargement as a consequence of lateral fusion and/or accumulation of more lipids generate curvature of the ER membrane. At this stage acyl-CoA synthetase 3 (ACSL3), an enzyme that provides acyl-CoA for LD formation, moves within the ER and becomes concentrated at emerging LD sites through its amphipathic α helices (*Kassan et al., 2013; Poppelreuther et al., 2012*). This enzyme is critical for LD expansion and maturation. Loss or overexpression of other ACSL family members does not affect LD biogenesis (*Kassan et al., 2013*), highlighting the importance of ACSL3-mediated local synthesis of acyl-CoA for LD expansion. ACSL3 belongs to the class I LD proteins (*Kory et al., 2016*), which are characterized by their presence in the ER without LDs and translocation to the LD surface during LD formation or after reconnection of LDs to the ER via membrane bridges (*Wilfling et al., 2013*). Most class I proteins have hydrophobic regions with hairpin-like structures (*Kory et al., 2016*). Emerging LDs expand and then are recognized by proteins such as perlipins. Perlipins are class II proteins containing amphipathic α helices or other hydrophobic domains that move from the cytosol to the LD surface (*Kory et al., 2016*).

Syntaxin 17 (Stx17) was originally characterized as a t-SNARE located in the smooth ER (*Steegmaier et al., 2000*). Recent studies by us and other groups demonstrated that Stx17 localizes to the mitochondria-ER interface, including the mitochondria-associated ER membrane (MAM) (Vance, 2014), and plays roles in mitochondrial division (*Arasaki et al., 2015*), the fusion of stress-induced mitochondrial-derived vesicles with lysosomes (*McLelland et al., 2016*), and autophagy in starved cells (*Diao et al., 2015; Hamasaki et al., 2013; Itakura et al., 2012; Takáts et al., 2013*). In mitochondrial division and autophagosome formation, Stx17 functions not as a SNARE, but as a scaffold at the MAM (*Arasaki et al., 2015; Hamasaki et al., 2013*). For these functions and the MAM localization, the C-terminal hairpin-like hydrophobic domain (CHD) and the subsequent cytoplasmic tail of Stx17, but not its SNARE domain, are important (*Arasaki et al., 2015*). The MAM is enriched in cholesterol and sphingolipids, thus resembling lipid raft-like structures (*Chipuk et al., 2012; Hayashi and Su, 2007; Sano et al., 2009*), although the ER membrane is very poor in cholesterol. The MAM has versatile functions (*Vance, 2014*), including the synthesis of neutral lipids as well as phospholipids (*Rusiñol, et al., 1994; Stone et al., 2009*).

In the present study we demonstrated that Stx17 is required for LD biogenesis. We found that Stx17 interacts with ACSL3, and facilitates its translocation from the ER to the surface of LDs.

## Results

### Stx17 is required for LD biogenesis

Although Stx17 is ubiquitously expressed, it is abundantly expressed in steroidogenic and hepatic cells (*Steegmaier et al., 2000*), both of which have large numbers of LDs. This and the MAM localization of Stx17 prompted us to examine the role of Stx17 in LD biogenesis. We first examined whether Stx17 is required for LD biogenesis by silencing the protein. We used two siRNAs (siRNA 440 and 194) that were able to effectively knockdown Stx17 (*Arasaki et al., 2015, and Figure 1A*), and found that the size and number of LDs were significantly reduced in hepatic cells (HepG2 and Huh7 cells) depleted of Stx17 (*Figure 1B, left two columns*). Silencing of Stx17 also inhibited oleic acid (OA)-induced LD biogenesis in HeLa cells (*Figure 1B, right column*). In accordance with the inhibition of LD formation, TAG synthesis was blocked in Stx17-silenced HeLa cells (*Figure 1C*). Of note is that Stx17 was not localized on the surface of LDs (*Figure 1B, top row*), suggesting that Stx17 does not directly participate in LD biogenesis, but rather has a regulatory role. To verify the physiological importance of Stx17 in LD biogenesis, we examined the effect of Stx17 silencing on the differentiation of 3T3-L1 preadipocytes into adipocytes. As differentiation progressed, the expression level of Stx17 markedly increased (*Figure 1-figure supplement 1A*). Silencing of Stx17 inhibited adipocyte differentiation, as demonstrated by a reduced increase in Oil-Red O staining (*Figure 1-figure supplement 1B,C*).

**Figure 1.**
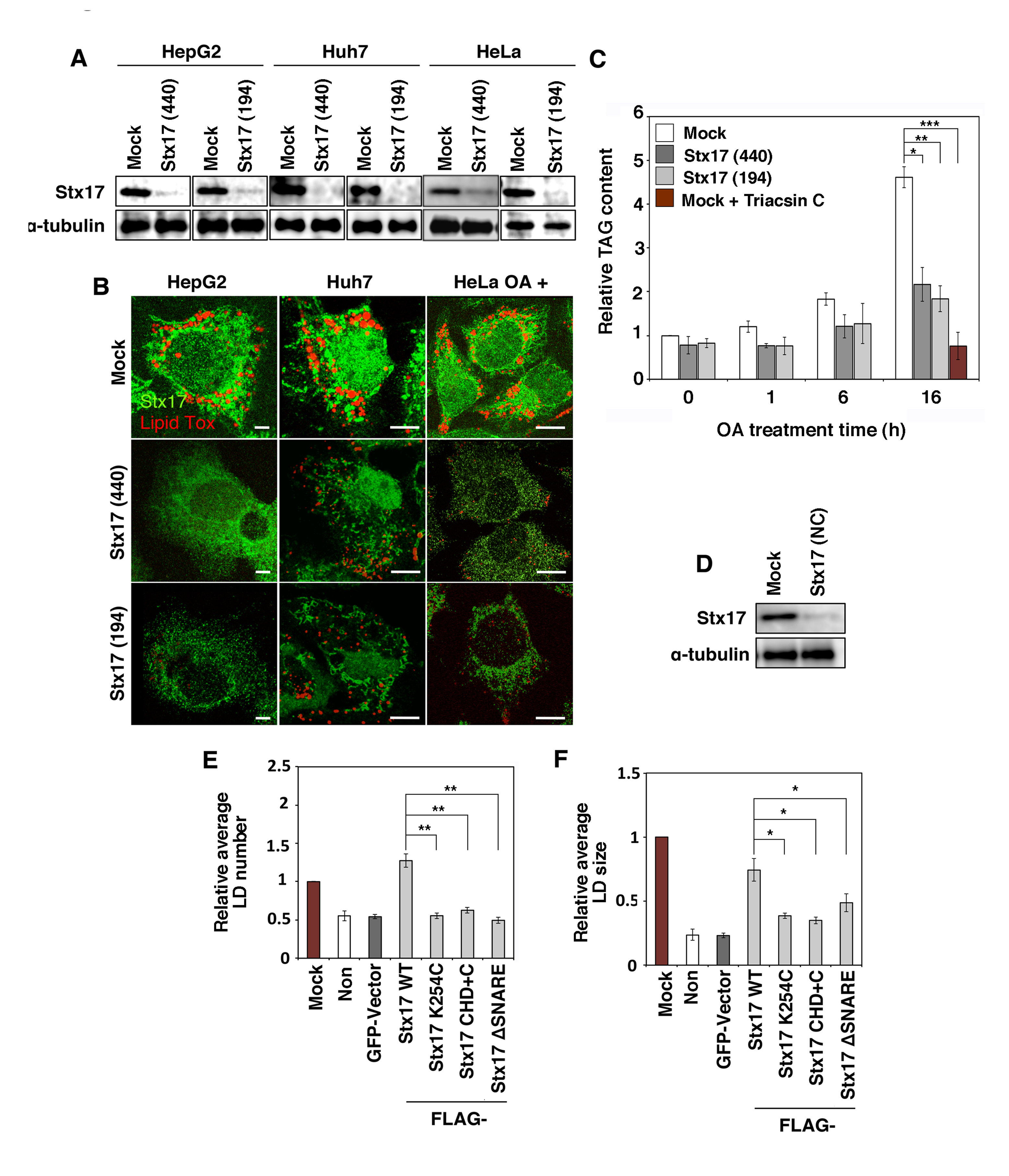
LD formation and TAG synthesis are impaired in Stx17-depleted cells. (**A,D**) HepG2, Huh7, and HeLa cells were mock-transfected or transfected with siRNA Stx17 (440), (194), or (NC) targeting the 3’ non-coding region of Stx17, and the protein amounts of Stx17 and α-tubulin were determined using their antibodies. (**B**) HepG2, Huh7, and HeLa cells were mock-transfected or transfected with siRNA Stx17 (440) or (194). At 72 hr after transfection, cells were fixed and stained with an anti-Stx17 antibody and Lipid Tox. For HeLa cells, OA was added at a final concentration of 150 μM at 56 hr after transfection of siRNA, followed by incubation for 16 hr. Bars, 5 μm. (**C**) HeLa cells were mock-transfected or transfected with siRNA Stx17 (440) or (194), treated with OA for the indicated times, and lysed, and then the amount of TAG was determined using an adipogenesis colorimetric/fluorometric assay kit. As a negative control, mock-treated HeLa cells were incubated with OA in the presence of 10 μM Triacsin C for 16 hr, and then the amount of TAG was determined. The bar graph shows the means ± SD (n = 3). (**E,F**) HeLa cells or HeLa cells expressing the indicated FLAG-tagged constructs were transfected with siRNA Stx17 (NC), treated with OA for 16 hr, fixed, and then immunostained with an anti-FLAG antibody and Lipid Tox. The graphs show the average number (**E**) and size (**F**) of LDs under each condition. Values are the means ± SD (n = 3).

To gain an insight into the mechanism by which Stx17 participates in LD biogenesis, we examined which domains of Stx17 are responsible for LD biogenesis. To address this, we performed rescue experiments using siRNA (Stx17 (NC)) that targets the 3’ non-coding region of Stx17 (*Figure 1D*). In Stx17-silenced cells, FLAG-tagged Stx17 wild-type showed restored size and number of LDs, excluding the possibility of an off-target effect of the siRNAs used (*Figure 1E,F*). We examined the ability of several Stx17 mutants (*Figure 1-figure supplement 1D*) to compensate for Stx17 depletion. No rescue was observed for Stx17 K254C in which Lys254 in the middle of the CHD was replaced by Cys, the CHD+C mutant, or the ΔSNARE mutant (*Figure 1E,F*), suggesting that both the SNARE domain and the CHD with the C-terminal cytoplasmic region, the latter of which is required for the MAM localization (*Arasaki et al., 2015*), are involved in LD biogenesis.

### Depletion of Stx17 causes aberrant distribution of ACSL3 on the LD surface

ACSL3, the most abundant acyl-CoA synthetase family member on LDs (*Brasaemle et al., 2004; Fujimoto et al., 2004*), redistributes from a microdomain of the ER to the LD surface during LD formation, and the enzymatic activity of ACSL3 plays a crucial role in LD expansion (*Fujimoto et al., 2007; Kassan et al., 2013*). On the other hand, depletion of ACSL4, the most abundant but non-LD-localized family member, does not affect LD formation, suggesting that the presence of ACSL on the LD surface is important for LD expansion (*Kassan et al., 2013*). Because TAG synthesis was markedly compromised in Stx17-depleted cells, we reasoned that Stx17 depletion might have caused ACSL3 dysfunction. Therefore, we examined the localization of ACSL3 in Stx17-depleted cells. As reported previously (*Kassan et al., 2013*), ACSL3 was detected on the surface of LDs exhibiting a circular distribution in mock-treated cells (*Figure 2A, left, upper row, B, top row*). In Stx17-silenced cells, on the other hand, ACSL3 exhibited a crescent-like distribution on the surface of LDs (*Figure 2A, left, lower row, B, second row*). Importantly, there was no difference in the circular distribution of Tip47/PLIN3, a LD-localized protein that redistributes from the cytosol, between mock- and Stx17-silenced cells (*Fig. 2A, right, B, third and bottom rows*). Quantification revealed that the recruitment of ACSL3 to the surface of LDs was specifically suppressed by Stx17 knockdown (*Figure 2C*).

**Figure 2.**
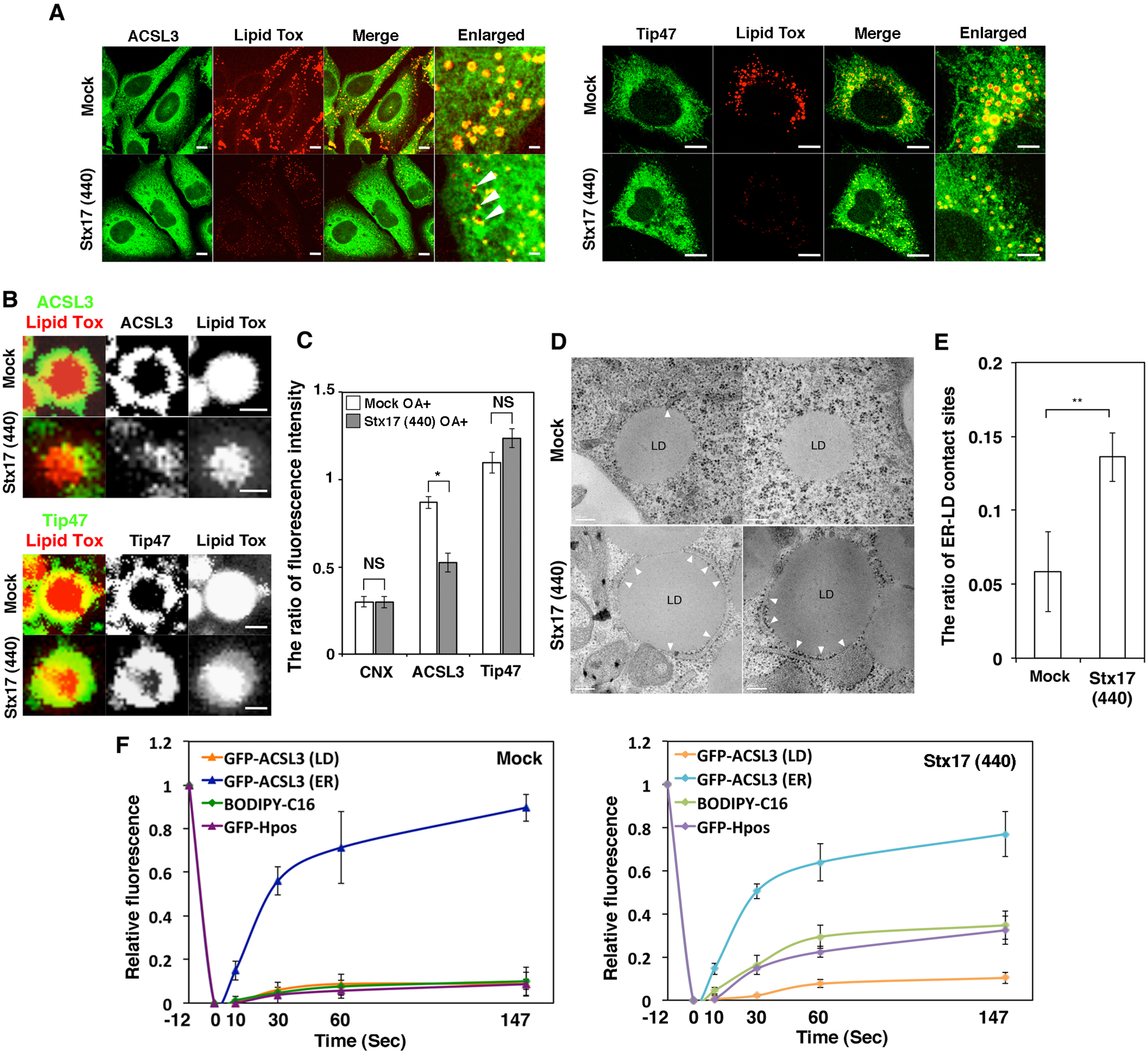
Aberrant distribution of ACSL3 on LDs in Stx17-silenced cells. (**A**) HeLa cells were mock-transfected or transfected with siRNA Stx17 (440), treated with OA for 16 hr, fixed, and then stained with Lipid Tox and an antibody against ACSL3 or Tip47. Bars in normal images, 5 μm, and in enlarged images, 1 μm. (**B**) Enlarged images of (**A**). Bars, 1 μm. (**C**) Quantitation of the data in (**B**). The fluorescence intensities of calnexin (CNX), ACSL3, and Tip47 surrounding LDs relative to that of Lipid Tox were plotted. The bar graph shows the means ± SEM (n = 3). (**D**) Electron micrographs of mock-treated HeLa cells (upper row) and Stx17-silenced cells (lower row) after 16 hr OA treatment. Bars, 200 nm. (**E**) Quantitation of the data in (**D**). The ratio of the area of ER-LD contact sites relative to the circumference of the LD surface was plotted. The bar graph shows the means ± SD (n = 5). More than 30 LDs were analyzed in each experiment. (**F**) Mock-treated or Stx17-silenced cells were transfected with one of the indicated plasmids or treated with BODIPY FL-C16 to visualize the surface or luminal domain of LDs. FRAP experiments were performed as described under Materials and methods. The graphs show the relative signal intensity after photobreaching. Values are the means ± SD (n = 3). The raw data are shown in *figure 2-figure supplement 1*.

To gain an additional morphological insight in Stx17-depleted cells, we performed electron microscopy. As shown in *Figure 2D,E*, the area of LDs that is in contact with the ER was significantly increased in Stx17-silenced cells. Because LDs bud from the ER and finally become detached, our electron microscopic data suggest that LDs remain immature and attached to the ER due to loss of Stx17. To directly test this, we performed FRAP experiments. In mock-treated cells, the signals of LD-localized BODIPY FL-C16, a membrane-permeable fatty acid that is efficiently incorporated into the triacylglycerol pool (*Rambold et al., 2015; Somwar et al., 2011*), as well as those of GFP-HPos, which redistributes from the ER to LDs during LD formation (*Kassan et al., 2013*), were not recovered after photobleaching (*Figure 2F, left, and Figure 2-figure supplement 1A*), consistent with the view that mature LDs become detached from the ER. On the other hand, significant recovery of the signals derived from these probes was observed after photobleaching of Stx17-silenced cells (*Figure 2F, right, and Figure 2-figure supplement 1B*), implying that LDs remained associated with the ER. Nevertheless, little recovery was observed for LD-localized GFP-ACSL3. In contrast to LD-localized GFP-ACSL3, an ER-localized GFP-ACSL3 fraction was rapidly recovered regardless of whether Stx17 was present or not (*Figure 2F and Figure 2-figure supplement 1A,B, bottom row*). These results suggest that Stx17 is a critical factor for the escort of GFP-ACSL3 from the ER to LDs.

### The GATE domain of ACSL3 is important for the interaction with the SNARE domain of Stx17

Because the redistribution of ACSL3 to LDs was suppressed in Stx17-silenced cells, Stx17 might regulate the localization of ACSL3 through protein-protein interaction. To test this possibility, we performed immunoprecipitation and proximity ligation assay (PLA). Significant amounts of GFP-tagged ACSL3 (*Figure 3A*) and endogenous ACSL3 (*Figure 3B*) were co-precipitated with FLAG-Stx17 wild-type and the K254C mutant, but not with the ΔSNARE mutant. Similar results were obtained for PLA (*Figure 3C and Figure 3-figure supplement 1A*), suggesting that Stx17 interacts with ACSL3 via its SNARE domain. We next examined which domains of ACSL3 are required for the interaction with Stx17. ACSL3 has transmembrane domain (TMD) and a GATE domain (*Figure 3D*), the latter of which is implicated in the control of access of the fatty acid substrate to the catalytic site of each of the two subunits (*Hisanaga et al., 2004; Soupene et al., 2010*). GFP-ACSL3 wild-type was found to decorate LDs (*Figure 3- figure supplement 1B, top row*). On the other hand, a dot-like distribution at the edge of LDs was observed for the ΔGATE mutant (*middle row*),and expression of the ΔTMD mutant reduced the number of LDs (*bottom row*). Binding experiments revealed that both the TMD and GATE domains of ACSL3 are required for the interaction with Stx17 (*Figure 3E*), although the ΔGATE mutant as well as the wild-type protein may be close to Stx17 in the absence of OA (*Figure 3F*). These findings combined with the fact that the SNARE domain of Stx17 is required for LD formation (*Figure 1E*,*F*) suggest that the interaction of ACSL3 with Stx17 is important for the redistribution of ACSL3 to LDs.

**Figure 3.**
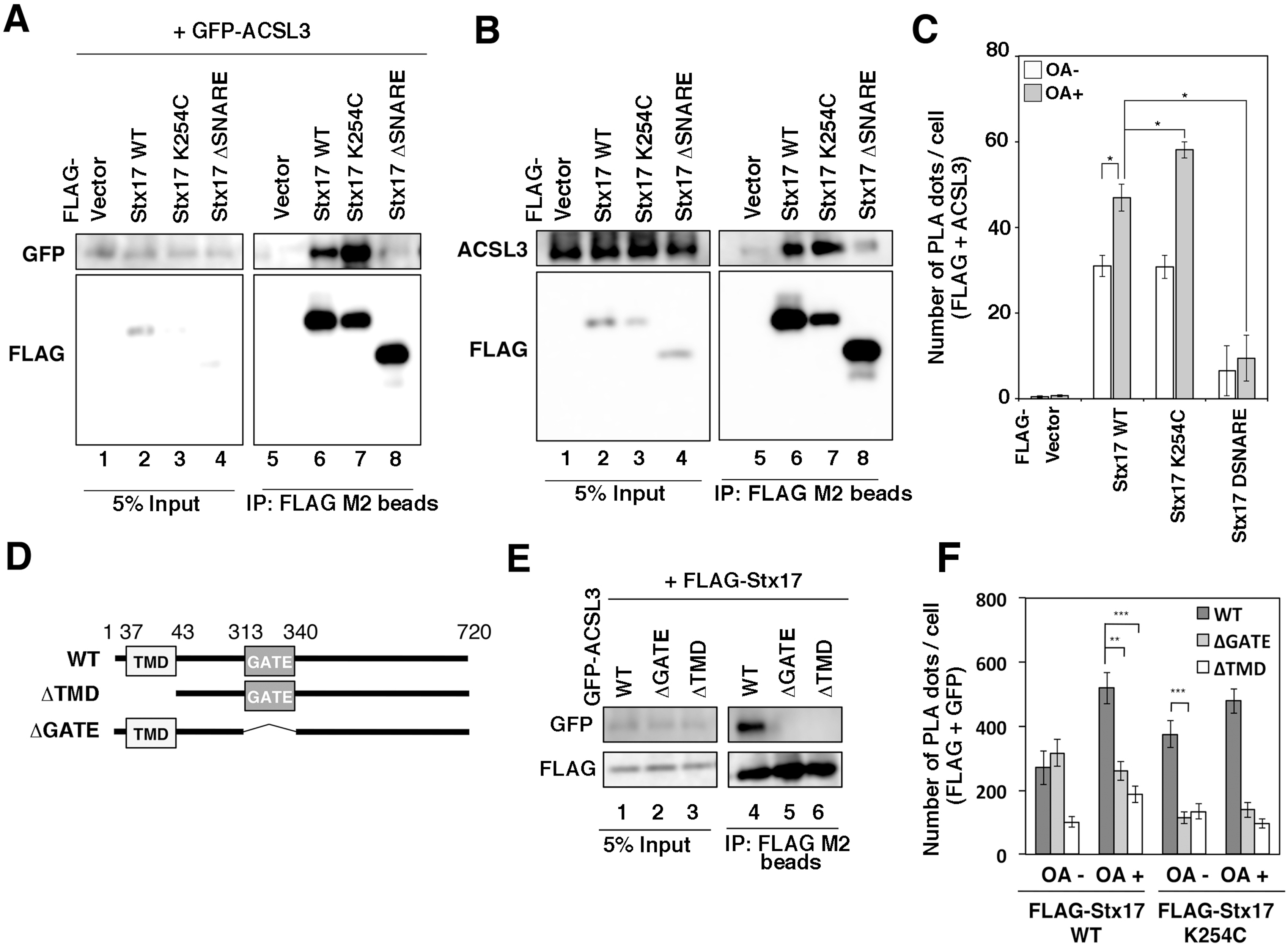
The SNARE domain of Stx17 and GATE domain of ACSL3 are indispensable for the interaction between Stx17 and ACSL3. (**A,B,E**) HeLa cells were transfected with the indicated constructs. At 24 hr after transfection, cell lysates were prepared and immunoprecipitated using anti-FLAG M2 beads. 5% input and the precipitated proteins were analyzed with the indicated antibodies. (**C**) HeLa cells were transfected with one of the indicated FLAG-Stx17 constructs and then incubated for 24 hr. Cells were incubated in the presence (dark gray bar) or absence (white bar) of OA for 16 hr and fixed, and PLA was performed using antibodies against FLAG and ACSL3. The bar graph shows the means ± SEM (n = 3). *Figure 3-figure supplement 1A*. (**D**) Schematic representation of ACSL3 and its deletion mutants. (F) HeLa cells stably expressing Stx17 wild-type or the K254C mutant were transfected with GFP-ACSL3 wild-type (dark gray bar), the ΔGATE mutant (light gray bar), or the ΔTMD mutant (white bar), and treated as described in (C). PLA was performed using antibodies against FLAG and GFP. The bar graph shows the means ± SEM (n = 3).

We next examined whether Stx17 regulates the localization of other enzymes that function in LD biogenesis. Lysophosphatidylcholine acyltransferase 1/2 (LPCAT1/2) are lipid synthesis enzymes that are related to the Lands cycle and relocate from the ER to LDs during LD maturation to regulate the size of LDs (*Moessinger et al., 2011, 2014*). Diacylglycerol acyltransferase 1/2 (DGAT1/2) produce TAG, and DGAT2 is known to localize to LDs as well as MAMs (*Kuerschner et al., 2008; Stone et al., 2009; Wilfling et al., 2013*). Among these enzymes, Stx17 interacts with LPCAT1 (*Figure 3 - figure supplement 1C,D*) and regulates its distribution (*figure supplement 1E*), whereas no significant translocation defect was observed for LPCAT2, the closest homologue of LPCAT1 among the LPCAT family proteins (Shindou and Shimizu, 2009). These findings suggest that Stx17 specifically regulates the redistribution of some enzymes to LDs through protein-protein interactions.

### SNAP23 localizes to the MAM

Given that Stx17 regulates the redistribution of ACSL3 from the ER to LDs likely through protein-protein interaction, we reasoned that other protein(s) might modulate this interaction. We focused on SNAP23 because previous studies revealed the involvement of this protein in lipid droplet formation and localization (*Boström et al., 2007; Jägerström et al. 2009*), and our interactome analysis identified SNAP23 as an Stx17-interacting protein (data not shown). We examined whether SNAP23, like Stx17, localizes to the MAM, in addition to the ER and mitochondria. As shown in *Figure 4A*, SNAP23 was found to be highly enriched in the MAM fraction. Of note is that a significant amount of ACSL3 was also recovered in the MAM fraction.

**Figure 4.**
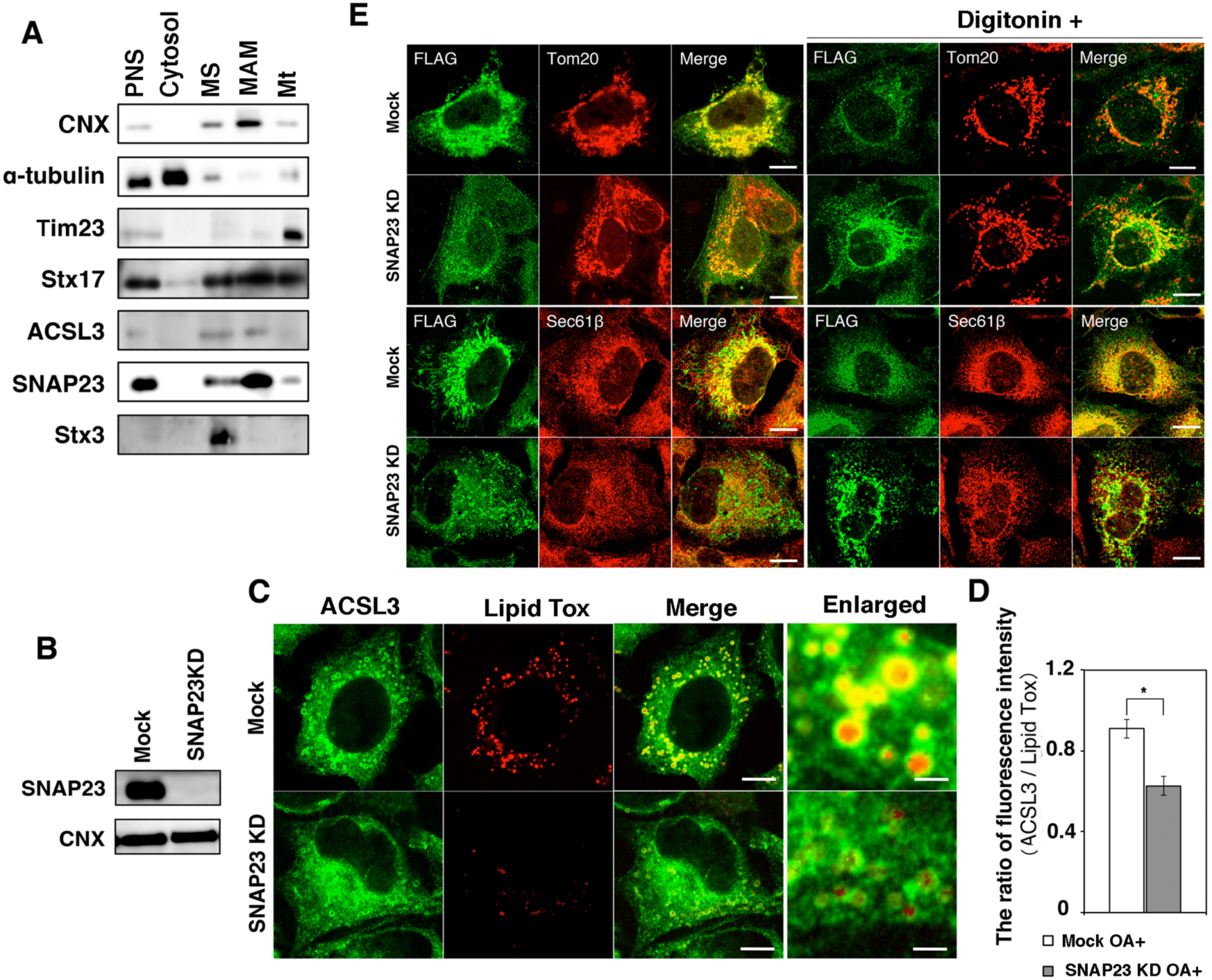
SNAP23 localizes to the MAM. (**A**) Subcellular fractionation of HeLa cell lysates was performed as described under Materials and methods. Equal amounts of proteins in individual fractions were subjected by SDS-PAGE and analyzed with the indicated antibodies. PNS, postnuclear supernatant; MS, microsomes; Mt, mitochondria. (**B**) HeLa cells were mock-transfected or transfected with siRNA targeting SNAP23, and protein levels were determined with antibodies against Stx17 and CNX. (**C**) Mock-treated and SNAP23-depleted cells were treated with OA for 16 hr, and then stained with an anti-ACSL3 antibody and Lipid Tox. Bars, 5 μm. (**D**) Quantitation of the data in (**C**). The graph shows the relative ACSL3 staining intensity surrounding LDs. Values are the means ± SEM (n = 3). (**E**) HeLa cells stably expressing FLAG-Stx17 were mock-transfected or transfected with siRNA targeting SNAP23. After 72 hr, cells were fixed without treatment (*left*) or treated with 30 μg/ηl digitonin for 5 min at room temperature (*right*), and then fixed, followed by immunostaining with antibodies against FLAG and Tom20 or Sec61β. Bars, 5 μm.

Silencing of SNAP23 (*Figure 4B*) markedly reduced the size of LDs, as previously reported (*Boström et al., 2007; Jägerström et al. 2009*), with a disrupted distribution of ACSL3 on LDs (*Figure 4C,D*), reminiscent of that observed in Stx17-silenced cells (*Figure 2A,B*). SNAP23 depletion caused a change in the distribution of Stx17 from a mitochondria-like pattern to a pattern with punctate structures (*Figure 4E, left, second and bottom rows*). Our previous study showed that the distribution of Stx17 is significantly perturbed upon treatment of cells with a low concentration (30 μg/ml) of digitonin (*Arasaki et al., 2015*), a reagent that can efficiently extract cholesterol (*Oliferenko et al., 1999*). This likely reflects the fact that the MAM is enriched in cholesterol and sphingolipids, thus resembling lipid raft-like structures (*Chipuk et al., 2012; Hayashi and Su, 2007; Sano et al., 2009*). We found that punctate Stx17-positive structures in SNAP23-depleted cells were relatively resistant to 30 μg/ml digitonin treatment (*Figure 4E, right, second and bottom rows*) compared to the mitochondria-like distribution of Stx17 in mock-treated cells (*top and third rows*). This suggests that Stx17 failed to gain access to raft-like structures in the absence of SNAP23.

### SNAP23 and ACSL3 bind exclusively to Stx17

We next determined the region of Stx17 responsible for the interaction with SNAP23 by immunoprecipitation (*Figure 5A*) and PLA (*Figure 5B*). As expected, the interaction of Stx17 with SNAP23 was abolished on deletion of the SNARE domain of Stx17 (*Figure 5A, lane 8*, B). Interestingly, the interaction was drastically reduced on mutation at Lys254 located in the middle of the CHD, even though the resultant mutant retained the SNARE domain (*Figure 5A, lane 7*,*B*).

**Figure 5.**
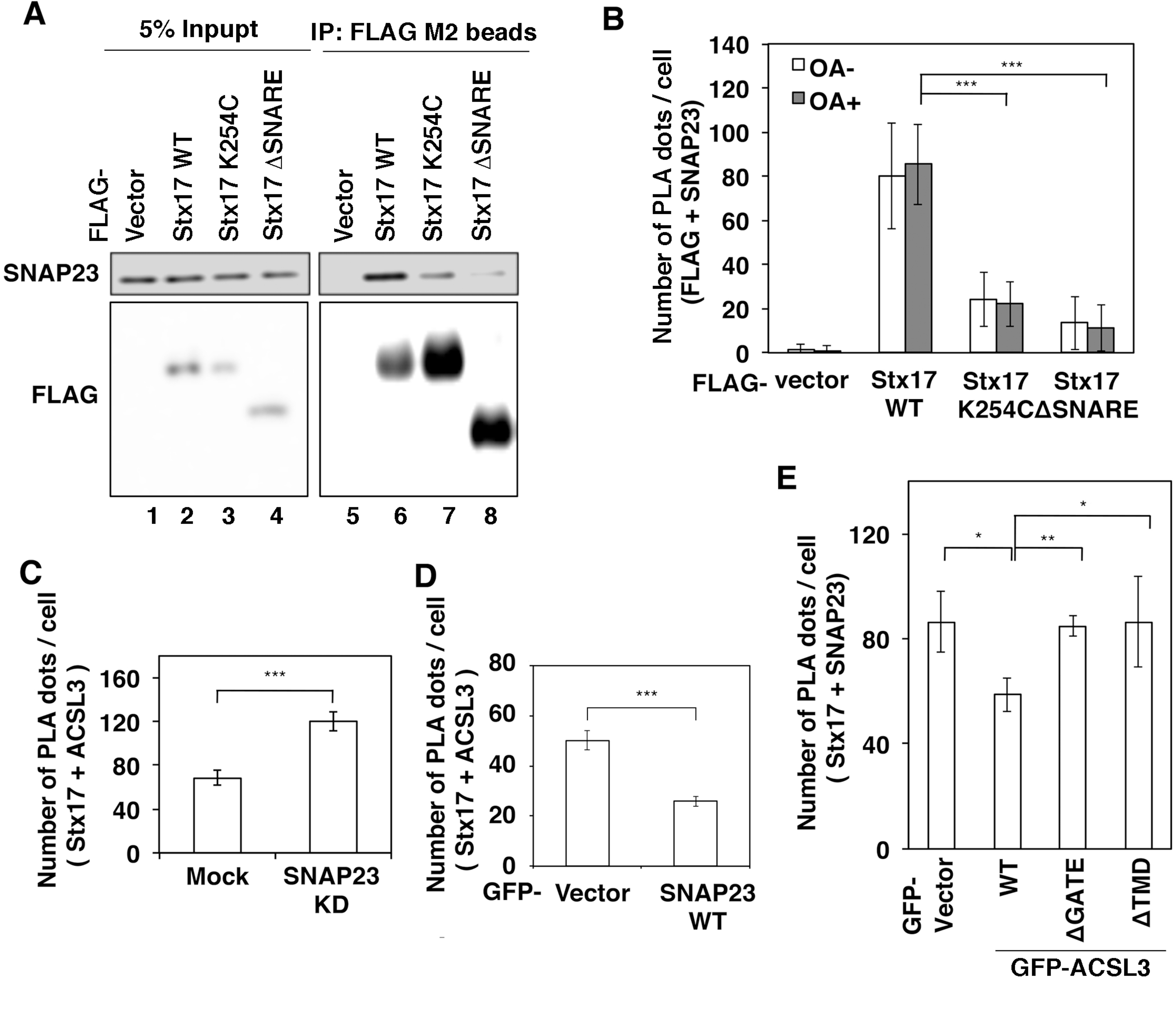
Exclusive binding of SNAP23 and ACSL3 to Stx17. (**A**) HeLa cells were transfected with one of the indicated FLAG-tagged constructs and then incubated for 24 hr. Cell lysates were prepared and immunoprecipitated using anti-FLAG M2 beads. 5% input and the precipitated proteins were analyzed with antibodies against SNAP23 and FLAG. (**B**) HeLa cells were transfected with one of the indicated FLAG-tagged constructs and then incubated for 24 hr. Cells were fixed, and PLA was performed using antibodies against FLAG and SNAP23. (**C,D**) HeLa cells were mock-transfected or transfected with siRNA targeting SNAP23 (**C**), or transfected with the GFP vector or GFP-SNAP23 wild-type (**D**). At 72 hr (siRNA transfection) or 24 hr (plasmid transfection), cells were incubated with OA for 16 hr. PLA was performed using antibodies against Stx17 and ACSL3. (**E**) HeLa cells were mock-transfected or transfected with one of the indicated GFP constructs. At 24 hr after transfection, cells were incubated with OA for 16 hr. PLA was performed using antibodies against Stx17 and SNAP23. (**B-E**) The bar graph shows the means ± SEM (n = 3).

Because both ACSL3 and SNAP23 interact with the SNARE domain of Stx17, we sought to determine whether they bind to Stx17 in a synergic or competitive manner. When SNAP23 was knocked down, the number of PLA dots representing the proximity between Stx17 and ACSL3 was significantly increased (*Figure 5C*), whereas ectopic expression of SNAP23 reduced the number of the PLA dots (*Figure 5D*). Furthermore, in cells expressing GFP-ACSL3 wild-type but not mutants lacking the ability to bind Stx17, i.e., the ΔGATE and ΔTMD mutants, the proximity of Stx17 to SNAP23 was disrupted (*Figure 5E*). These findings suggest that SNAP23 and ACSL3 compete for Stx17 binding.

### The MAM, but not tethering between the MAM and mitochondria, is important for LD formation

Next, we examined the effect of depletion of PACS-2, a multifunctional sorting protein that is required for maintaining MAM integrity (*Myhill et al., 2008; Simmen et al., 2005*), and mitofusin 2 (Mfn2), a key tether for ER-mitochondria (*Naon et al., 2016*). As shown in *Figure 6A*, Mfn2 depletion did not affect OA-induced LD formation, whereas PACS-2 was found to be required for LD formation. Consistent with these findings, the proximity signal for FLAG-Stx17 and ACSL3 was reduced upon depletion of PACS-2, but not Mfn2, and OA increased the signal (*Figure 6B*).

**Figure 6.**
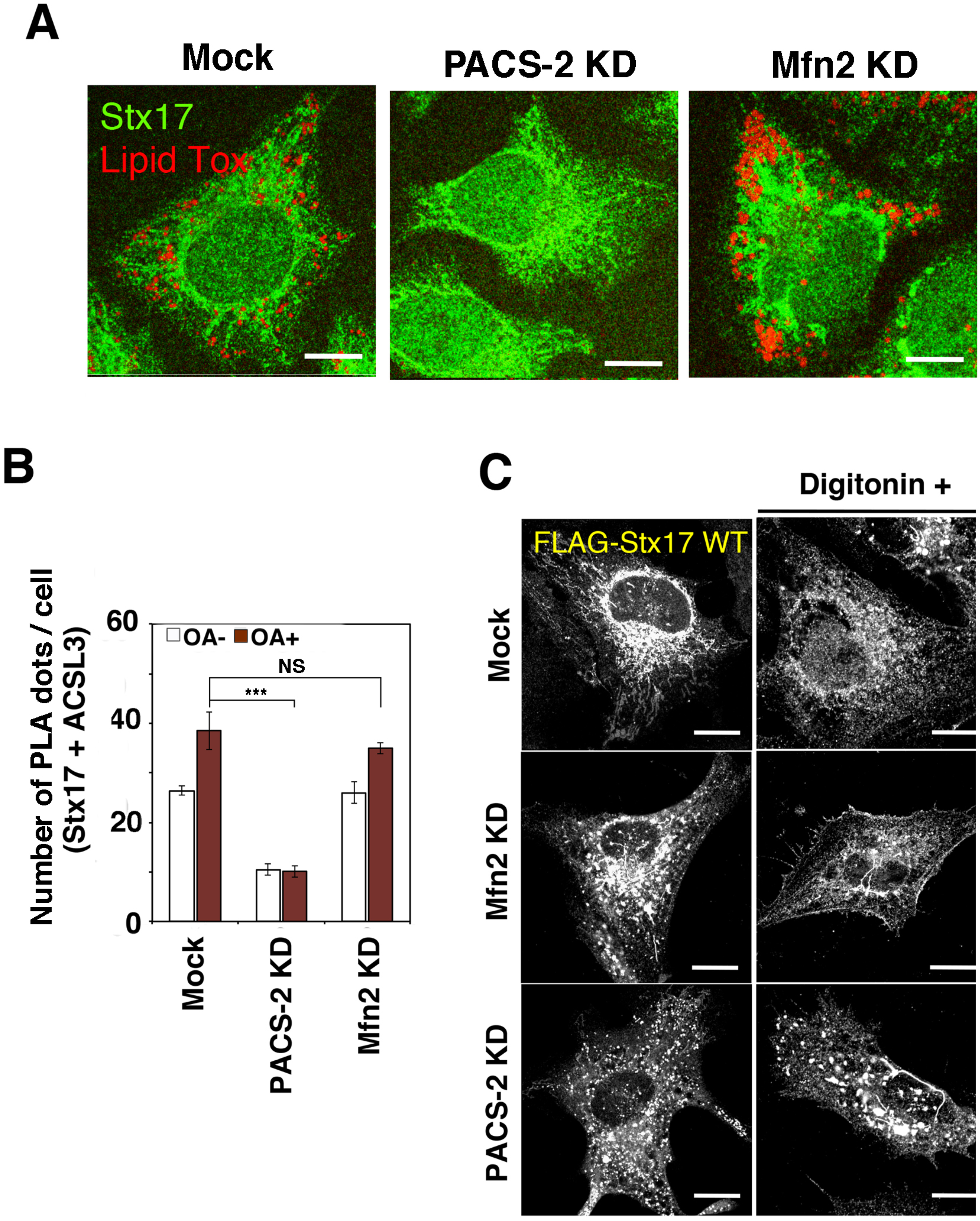
MAM is important for LD formation and the Stx17-ACSL3 interaction. (**A**) HeLa cells were mock-transfected or transfected with siRNA targeting PACS-2 or Mfn2. At 72 hr after transfection, cells were fixed and stained with an anti-Stx17 antibody and Lipid Tox. Bars, 5 μm. (**B**) Alternatively, PLA was performed using antibodies against Stx17 and ACSL3. The bar graph shows the means ± SEM (n = 3). (**C**) HeLa cells stably expressing FLAG-Stx17 were mock-transfected or transfected with siRNA siRNA targeting Mfn2 or PACS-2, treated with DMSO or 30 μg/ml digitonin for 5 min at room temperature, fixed, and then immunostained with an anti-FLAG antibody. Bars, 5 μm.

We assessed the Stx17 milieu in cells depleted of PACS-2 or Mfn2 by means of digitonin sensitivity. Although some punctate Stx17-positive structures were observed in Mfn2-silenced cells, they were mostly abolished by digitonin treatment (*Figure 6C, middle row*). Similar to those observed in SNAP23-depleted cells, on the other hand, prominent punctate Stx17-positive structures were resistant to digitonin treatment (*bottom row*), suggesting that Stx17 fails to localize to raft-like structures in cells depleted of PACS-2 as well as SNAP23. Alternatively, raft-like structures might have been disrupted upon depletion of PACS-2.

### Transient binding of Stx17 to ACSL3 during LD formation

Given that SNAP23 and ACSL3 compete for Stx17 binding, one attractive hypothesis for the regulation of the redistribution of ACSL3 from the ER to nascent LDs due to Stx17 and SNAP23 is that at the onset of LD formation ACSL3 first binds to Stx17, and then SNAP23 competitively binds to Stx17, releasing ACSL3 from Stx17 to allow its redistribution to the LD surface. To test this hypothesis, we monitored the change of the binding partner during LD maturation. Immunoprecipitation (*Figure 7A*) and PLA (*Figure 7B*) revealed that the binding of ACSL3 to Stx17 was augmented at 1-hr incubation with OA and then decreased, whereas the binding of Stx17 to SNAP23 increased up to 6 hr.

**Figure 7.**
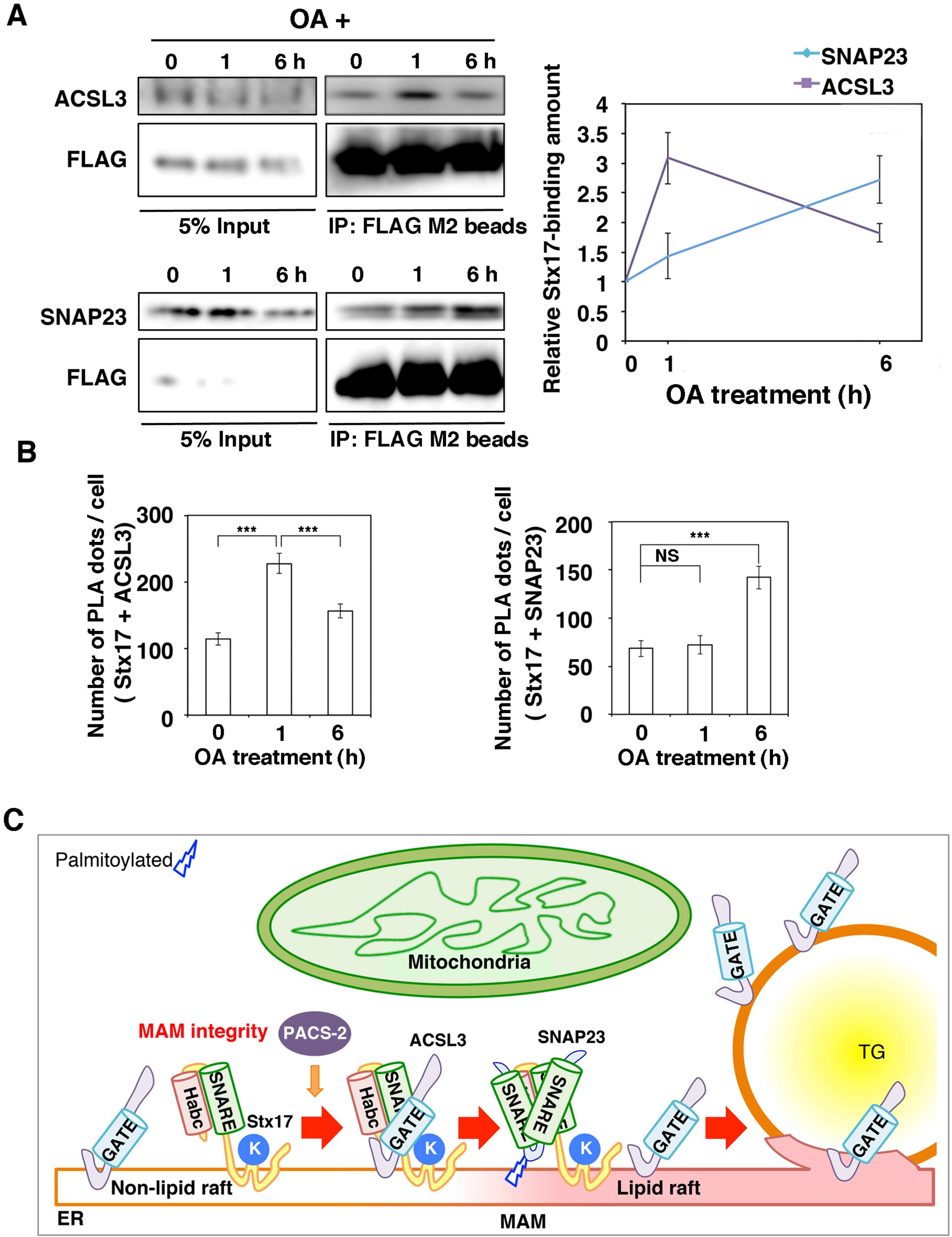
Stx17 changes the binding partner from ACSL3 to SNAP23 during LD maturation. (**A**) HeLa cells stably expressing FLAG-Stx17 were incubated with OA for the indicated times. Cell lysates were prepared and immunoprecipitated using anti-FLAG M2 beads. 5% input and the precipitated proteins were analyzed with the indicated antibodies. The graph on the right shows the relative band intensity. Values are the means ± SD (n = 3). (**B**) HeLa cells were treated with OA for the indicated times, and PLA was performed using antibodies against Stx17 and ACSL3 or SNAP23. The bar graphs show the means ± SEM (n = 3). (**C**) Working model of the function of Stx17 in LD maturation. For details, see Discussion.

## Discussion

The protein and lipid compositions of LDs vary not only with the cell type but also in the degrees of LD maturation and consumption in individual cell types (*Ohsaki et al., 2014; Pol et al., 2014; Thiam and Beller, 2017*). The importance of the recruitment of proteins to LDs in LD biogenesis is highlighted by the finding that depletion of LD-localized ACSL3 impaired LD formation, whereas depletion of the most abundant but non-LD-localized member of the ACSL family, ACSL4, had no impact on LD biogenesis (*Kassan et al., 2013*), suggesting that on site supply of acetyl-CoA is necessary for the growth of LDs. Therefore, how proteins selectively bind to the LD surface, which is surrounded by a phospholipid monolayer, in a maturation stage-dependent manner is one of the most key questions to be addressed in LD biogenesis. In this study, we demonstrated that Stx17, a SNARE protein localized in the MAM, in conjunction with another MAM protein, SNAP23, regulates the translocation of ACSL3 and perhaps LPCAT1 from the ER to nascent LDs. On the other hand, the distribution of some LD-localized proteins such as DGAT2 and Tip47/PLIN3, which translocate from the ER and cytosol to LDs, respectively, was found to be independent of Stx17. These findings suggest that Stx17 selectively interacts with and regulates the localization of a subset of proteins for LD biogenesis.

We found that the SNARE domain and Lys254 in the middle of the CHD of Stx17 are important for LD biogenesis. The SNARE domain of Stx17 was found to be responsible for the interaction with ACSL3 and SNAP23. For binding to SNAP23, not only the SNARE domain of Stx17, but also Lys254, which is a critical residue for MAM localization (*Arasaki et al., 2015*), is important. In the absence of SNAP23, Stx17 localization was relatively resistant to digitonin treatment. The resistibility of Stx17 to digitonin was also acquired on loss of PACS-2, a MAM organizer (*Myhill et al., 2008; Simmen et al., 2005*). Based on the time course experiment results (*Figure 7A,B*) and other data, we envisage that the interaction of Stx17 and ACSL3 begins at non-raft like structures at the onset of LD formation (*Figure 7C*), and then ACSL3 is released from Stx17 due to the binding of SNAP23 to Stx17 at digitonin-sensitive sites, likely lipid raft-like structures. SNAP23 is known to be palmitoylated at several Cys residues located in the middle of the protein (*Vogel and Roche, 1999*), and this modification is required for the membrane attachment and/or raft association of SNAP23 (*Salaün et al., 2005*). Palmitoylation is a signal for targeting of proteins to raft structures (*Salaun et al., 2010*) and the MAM (*Lynes et al., 2012*). Finally, ACSL3 enters the phospholipid monolayer surrounding LDs, where the mobility of ACSL3 is highly restricted compared to in the ER.

Although the SNARE domain of Stx17 is important for LD biogenesis, we do not favor the idea that Stx17 participates in the formation of bridges that connect the ER and nascent LDs or, as proposed previously for Stx5 (*Boström et al, 2007*), LD fusion. After initial formation, LDs retain functional connection with the ER through membrane bridges (*Jacquier et al., 2011; Wilfling et al., 2013*), and a subset of proteins (class I proteins: *Kory et al., 2016*) translocate to LDs likely through these bridges (*Ingelmo-Torres et al., 2009; Wilfling et al., 2013; Zehmer et al., 2008*). FRAP experiments after maturation of LDs revealed that LDs are not connected with the ER in mock-treated cells, whereas in Stx17-depleted cells LDs are still in contact with the ER, as shown by the recovery of fluorescence of BODIPY FL-C16 and GFP-HPos. This is in line with the electron microscopic data revealing the close vicinity of small LDs to the ER membrane. Our idea that Stx17 is not involved in the formation of ER-LD bridges is supported by the finding that knockdown of Sec22b, a cognate v-SNARE for Stx17 (Steegmaier et al., 2000), does not affect LD formation (*data not shown*).

A recent study revealed that seipin, a transcript of the BSCL2 gene whose mutation causes severe congenital lipodystrophy (*Magré et al., 2001*), regulates the translocation of ACSL3 and lipids into LDs (*Salo et al., 2016*). Another recent study revealed that seipin suppresses the activity of glycerol-3-phosphate acyltransferase to prevent the formation of excess phosphatidic acid (*Pagac et al., 2016*). Both proteins are key enzymes for LD formation. Seipin and its yeast homologue, Fld1, localize to ER-LD contact sites (*Szymanski et al., 2007; Wang et al., 2016*), and loss of Fld1 or its partner, Ldb16, yields LDs with aberrant lipid and protein compositions (*Fei et al, 2011; Wang et al., 2014; Cartwright et al., 2015; Grippa et al., 2015*). Although seipin/Fld1 is conserved, Ldb16 is not present in mammals. Wang et al. (2014) postulated that Fld1 and Ldb16 in yeast might have evolved into a single molecule, seipin, in mammals. Our present observations revealed another protein that regulates the translocation of key enzymes from the ER to nascent LDs. Different from Fld1, Stx17 is not localized in ER-LD contact sites nor conserved in yeast. However, Stx17 is one of the Last Eukaryotic Common Ancestors (LECAs) and present in diverse eukaryotic organisms, but was also lost in multiple lineages (*Arasaki et al., 2015*). In organisms lacking Stx17 the role of Stx17 in LD formation may be assumed by another protein. Of note is that despite the absence of an obvious homologue in yeast, Stx17 plays pivotal roles at the initial and late stages in autophagy, a well-conserved and important process in eukaryotes (*Hamasaki et al., 2013; Itakura et al., 2012*).

## Materials and methods

### Chemicals and antibodies

Triacsin C, digitonin, and OA were obtained from Enzo Life Sciences, Wako Chemicals, and Sigma-Aldrich, respectively. Lipid Tox and BODIPY FL-C16 were obtained from Thermo Fisher Scientific. The following antibodies were purchased from Sigma-Aldrich: monoclonal FLAG (No. F3165) and polyclonal FLAG (No. F7425). The following antibodies were obtained from BD Bioscience Pharmingen: CNX (No. 610523), and Tom20 (No. 612278). The following antibodies were purchased from Proteintech: PACS-2 (No. 19508-1-AP), polyclonal SNAP23 (No. 10825-1-AP), and SNAP29 (No. 12704-1-AP). The following antibody was purchased from Santa Cruz Biotechnology: monoclonal SNAP23 (D-11: sc-374215). The following antibodies were from the indicated sources: HA (Sigma-Aldrich, H6908), Mfn2 (Abcam, ab56889), Alexa Fluor® 488 and 594 goat anti-mouse and -rabbit antibodies (Thermo Fisher Scientific, No. A-11001, A-11005, A-11008, and A-11012), and Sec61β (EMD Millipore, No. 07-205). An antibody against Stx17 was prepared as described previously (*Arasaki et al., 2015*) and used for immunofluorescence analysis. For immunoblotting, an anti-Stx17 antibody (Sigma-Aldrich; No. HPA001204) was used.

### Cell culture

293T, HepG2, Huh7, and 3T3-L1 cells were grown in DMEM supplemented with 50 IU/ml penicillin, 50 μg/ml streptomycin, and 10% fetal calf serum. HeLa cells (RIKEN, RCB0007) were cultured in α-MEM supplemented with the same materials plus 2 mM L-glutamine. Stable transfectants were prepared as described previously (*Arasaki et al., 2015*). For the induction of LDs in HeLa cells, OA was added at a final concentration of 150 μM.

### Plasmids and transfection

Plasmids encoding human Stx17 full-length and its derivatives were described previously (*Arasaki et al., 2015*). The plasmids for GFP-ACSL3, FLAG-DGAT1/2, and 3x-HA-LPCAT1/2 were gifts from Dr. Albert Pol (IDIBAPS), Dr. Robert V. Farese (Harvard University), and Dr. Christoph Thiele (University of Bonn), respectively. Deletion mutants of ACSL3 were constructed by inverse PCR. Transfection was carried out using LipofectAMINE Plus, LipofectAMINE2000 (Thermo Fisher Scientific), or PEI (*Longo et al., 2013*) according to the manufacturer’s protocol.

### RNA interference

The following siRNAs were used:

Stx17 (440) : 5’-GGUAGUUCUCAGAGUUUGAUU-3’

Stx17 (194) : 5 ’-CGAUCCAAUAUCCGAGAAAUU-3 ’

Stx17 (NC) : 5’-GGAAAUUAAUGAUGUAAGA-3 ’

Stx17 (421) : 5 ’-CACACUGGGGAGGCUGAAGCU-3 ’

Mfn2: 5 ’-AGAGGGCCUUCAAGCGCCA-3’

PACS2: 5’-AACACGCCCGUGCCCAUGAAC-3’

SNAP23: 5’-CAUUAAACGCAUAACUAAU-3’

SNAP29: 5’-UAUCAUCCAGCUUUCUAAGGUUUGG-3’

siRNAs were purchased from Japan Bio Services. HeLa, HepG2, Huh7, and 3T3-L1 cells were grown on 35-mm dishes, and siRNAs were transfected at a final concentration of 200 nM using Oligofectamine, LipofectAMINE2000, or LipofectAMINE2000 RNAi Max (Thermo Fisher Scientific) according to the manufacturer’s protocol.

### Immunoprecipitation

HeLa cells expressing FLAG-tagged proteins were lysed in lysis buffer (20 mM Hepes-KOH (pH 7.2), 150 mM KCl, 2 mM EDTA, 1 mM dithiothreitol, 1 μg/ml leupeptin, 1 μM pepstatin A, 2 μg/ml aprotinin, and 1 mM phenylmethylsulfonyl fluoride) containing 1% Triton X-100 or 10 mg/ml digitonin. After centrifugation, the supernatants were collected and immunoprecipitated with anti-FLAG M2 affinity beads (Sigma-Aldrich). The precipitated proteins were eluted with SDS sample buffer, and then analyzed by immunoblotting. Experiments were repeated two or three times with similar results.

### Immunofluorescence microscopy

For immunofluorescence microscopy, cells were fixed with 4% paraformaldehyde for 20 min at room temperature or ice-cold methanol at −20°C, and then observed under an Olympus Fluoview 300 or 1000 laser scanning microscope. Unless specifically stated, shown in figures are representative images of at least three independent experiments. To determine the fluorescence intensity ratio (*Figures 2C and 4D*), immunofluorescence images obtained were analyzed using ImageJ software (NIH).

### Electron microscopy

Cells were fixed with a mixture of 2% formaldehyde and 2.5% glutaraldehyde in 0.1 M sodium cacodylate buffer (pH 7.4) for 2 hr, and then prepared for conventional observation as described previously (*Ohsaki et al., 2016*). ER-LD contact length and the circumference of each LD were determined with ImageJ software. ER-LD contact sites were discerned when the distance between the ER membrane and the surface of an LD was 20 nm or less.

### FRAP

FRAP experiments were performed with an Olympus Fluoview 1000 laser scanning microscope equipped with a stage-top incubator (37°C, 5% CO_2_). To monitor LDs during FRAP experiments, cells expressing GFP constructs or labeled with BODIPY FL-C16 were incubated with Lipid Tox in Opti-MEM supplemented with 10% fetal calf serum before photobleaching. The minimum region defined with the Olympus Fluoview 1000 laser scanning microscope was photobleached using a 488-nm laser at 100% laser power for 2 s. After photobleaching, images were obtained at 0.5 s intervals. In each experiment 30 cells were used, and more than 3 LDs in each cell were bleached. Experiments were repeated three times,

### Differentiation of 3T3-L1 cells and measurement of intensity of Oil Red O staining

Differentiation was induced using an AdipoInducer Reagent (for animal cell) kit (Takara Bio, MK429) according to the manufacturer’s protocol. Briefly, 3T3-L1 cells were mock-transfected or transfected with siRNA Stx17 (421). At 72 hr after transfection, cells were incubated with DMEM supplemented with 50 IU/ml penicillin, 50 μg/ml streptomycin, and 10% fetal calf serum, plus 10 μg/ml insulin solution, 2.5 μM dexamethasone, and 0.5 mM 3-isobutyl-1-methylxanthine, for 48 hr. The medium was replaced with DMEM containing 10 μg/m insulin, and then incubated for ∼9 days. For Oil Red O staining, cells were fixed, washed with 60% isopropanol for 1 min, and then incubated with a 60% Oil Red O solution (99% isopropanol/0.3% Oil Red O) for 15 min at 37°C. After washing with 60% isopropanol and drying, Oil Red O was dissolved in 100% isopropanol and the OD at 500 nm was measured.

### PLA

PLA was conducted using a PLA kit (Sigma-Aldrich) according to the manufacturer’s protocol. Thirty cells were analyzed in each assay. PLA dots were identified using the “analyze particle” program in the ImageJ software. Randomly, 30 cells were selected and the number of PLA dots was measured in each sample, and the experiments were repeated three times.

### Subcellular fractionation

Subcellular fractionation was performed as described previously (*Arasaki et al., 2015*). Experiments were repeated two or three times with similar results.

### Digitonin treatment

Digitonin was freshly dissolved in DMSO before use. Cells were incubated for 5 min at room temperature with 30 μg/ml digitonin in KHM buffer (25 mM Hepes (pH 7.2), 125 mM potassium acetate, 2.5 mM magnesium acetate, 1 mM dithiothreitol, and 1 mg/ml glucose).

### Statistical analyses

The results were averaged, expressed as the mean ± SD or SEM, and analyzed using a paired Student’s t test. The p values are indicated by asterisks in the figures with the following notations: *p ≤ 0.05; **p ≤ 0.01; ***p ≤ 0.001. NS, not significant.

## Acknowledgements

We thank Dr. Albert Pol, Dr. Christoph Thiele, Dr. Robert V. Farese, and the Gladstone Institutes, UCSF, for the generous gifts of plasmids. We are indebted to Ms. Risako Yoshida, Mr. Toshiki Amemiya, and Ms. Miwa Yokoyama for their technical assistance. This work was supported in part by Grants-in-Aid for Scientific Research, #25291029 and #26650066 (to M.T.), and #26111520 and #26713016 (to K.A.), and the MEXT-Supported Program for the Strategic Research Foundation at Private Universities (to M.T., and K.A.) from the Ministry of Education, Culture, Sports, Science and Technology of Japan.

## Author contributions

HK, Conception and design, Acquisition of data, Analysis and interpretation of data, Drafting; KA, Conception and design, Acquisition of data, Analysis and interpretation of data, Drafting; MT, Conception and design, Drafting; UO, Acquisition of data, Analysis and interpretation of data; TF, Analysis and interpretation of data.

---

**Figure 1-source data 1.** Values, statistics, and exact p values for Figure 1C,E,F.

**Figure 1-figure supplement 1.**
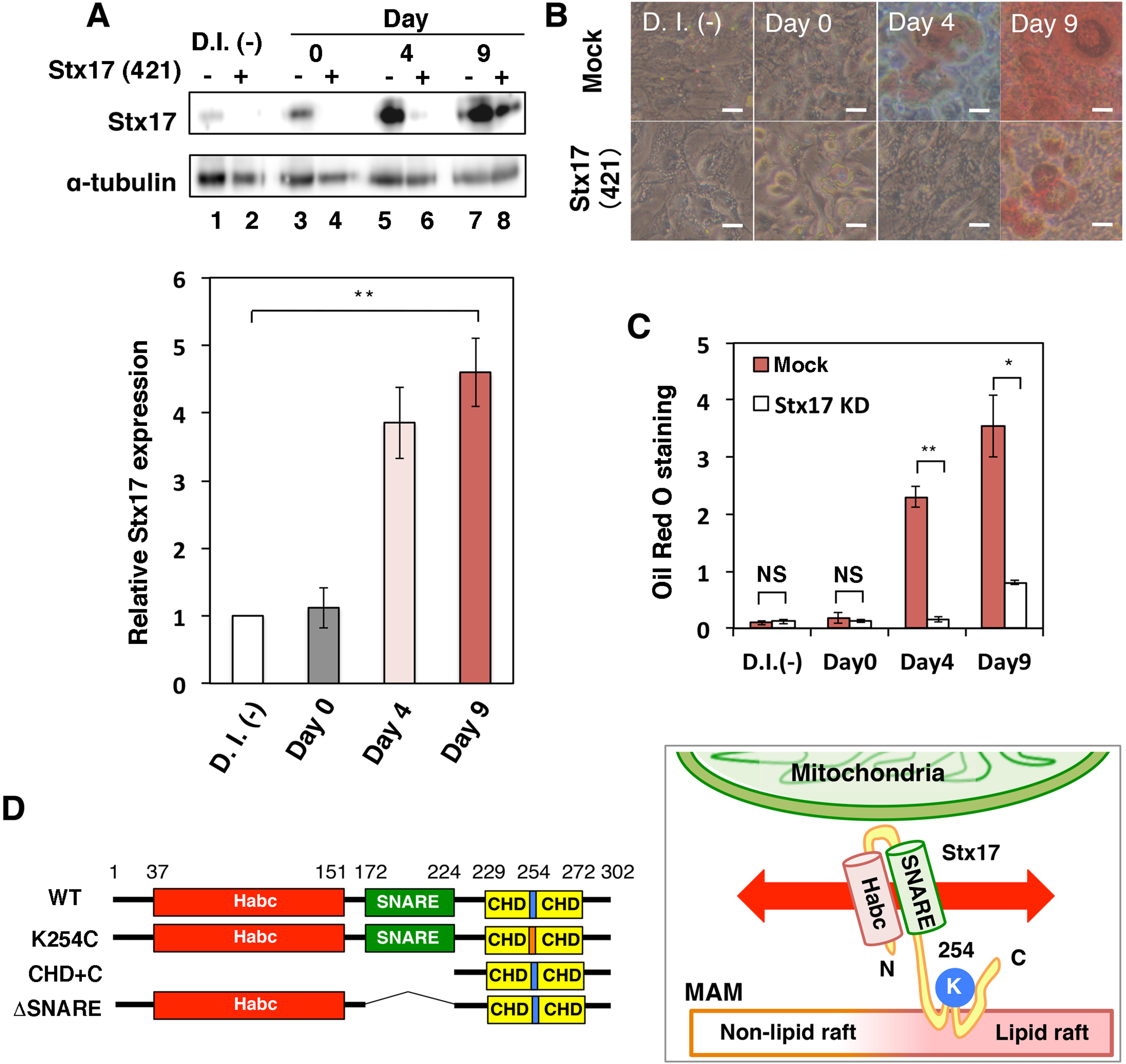
Silencing of Stx17 suppresses differentiation of 3T3-L1 preadipocytes. (A) 3T3-L1 cells were mock-transfected or transfected with siRNA targeting mouse Stx17 (siRNA (421)), and then differentiation was induced with an AdipoInducer Reagent. Protein expression levels were determined by immunoblotting using the indicated antibodies. The graph at the bottom shows the intensity of the Stx17 protein band relative to that of α-tubulin. D.I. (-) denotes no differentiation induction. The bar graph shows the means ± SD (n = 3). (**B**) Oil Red O staining of 3T3-L1 cells during differentiation. Bars, 5 μm. (**C**) Quantitation of Oil Red O staining. The bar graph shows the means ± SD (n = 3). (**D**) Distribution and schematic representation of Stx17 wild-type and its derivatives.

**Figure 1-source data 2.** Values, statistics, and exact p values for Figure 1-**f**igure supplement 1A,C.

**Figure 2-source data 1.** Values, statistics, and exact p values for Figure 2C,E,F.

**Figure 2-figure supplement 1.**
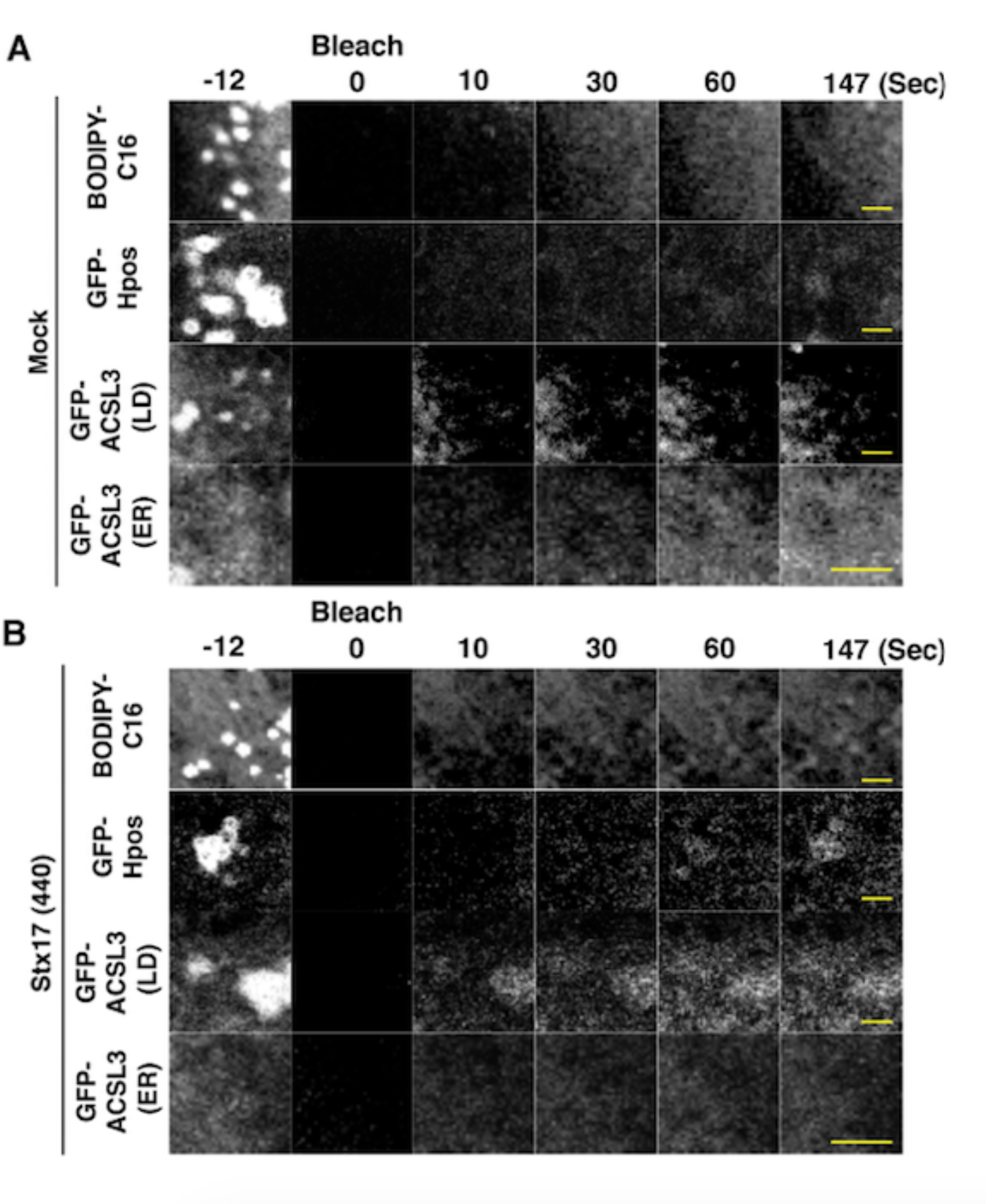
LDs are close to the ER in Stx17-silenced cells. The images in FRAP experiments. (**A**) Mock-treated cells and (**B**) Stx17-silenced cells. Bars, 5 μm.

**Figure 3-source data 1.** Values, statistics, and exact p values for Figure 3C,F.

**Figure 3-figure supplement 1.**
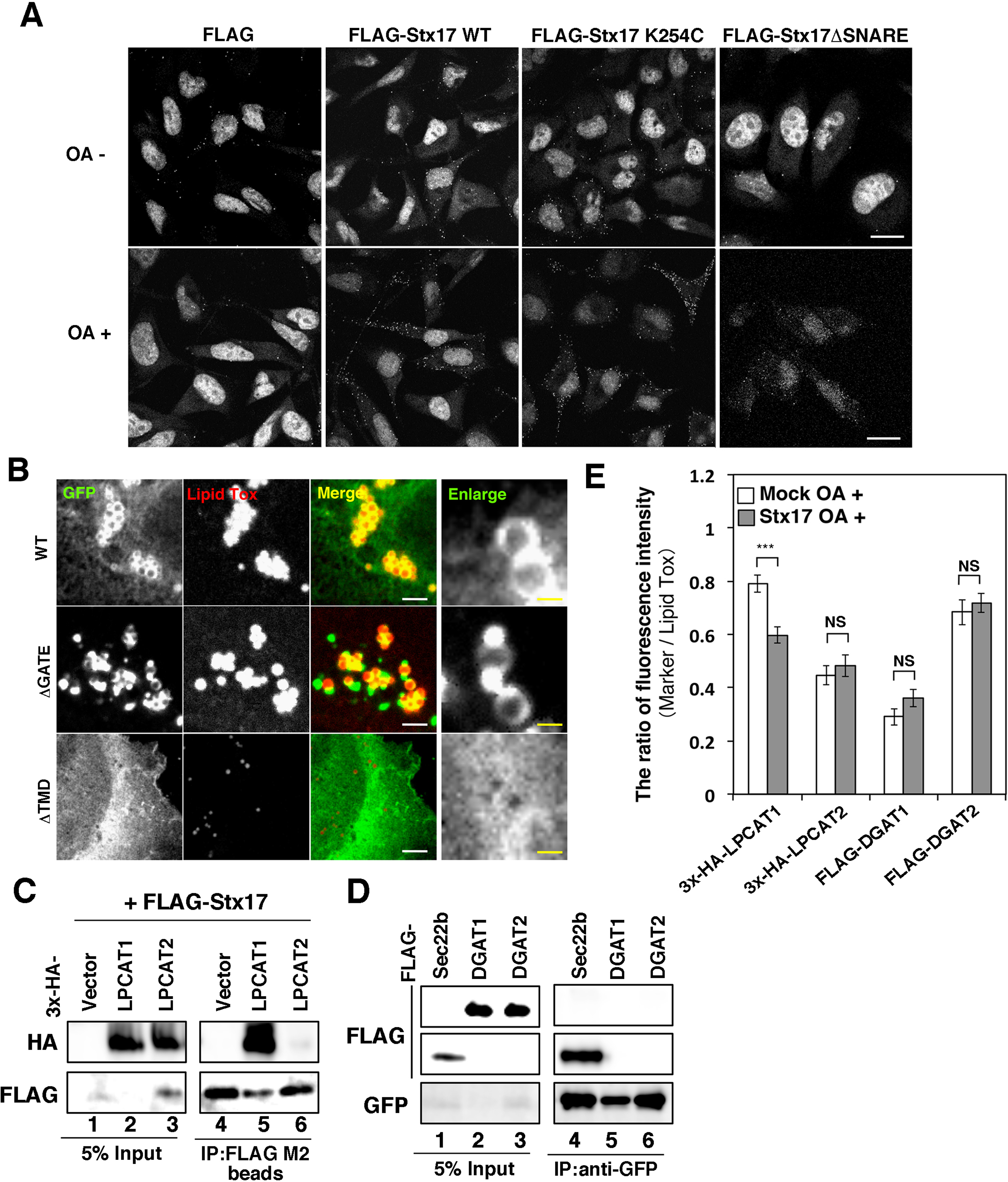
Localization of ACSL3 and LD-related proteins. (**A**) The PLA images related to *Figure 3C*. The fluorescence dots represent the proximity between Stx17 constructs and ACSL3. Bars, 5 μm. (**B**) HeLa cells were transfected with one of the indicated GFP-ACSL3 constructs. At 24 hr after transfection, cells were treated with OA for 16 hr, fixed, and then stained with Lipid Tox. (**C**) HeLa cells were transfected with FLAG-Stx17 and one of the indicated 3x-HA-tagged constructs. At 24 hr after transfection, cell lysates were prepared and immunoprecipitated using anti-FLAG M2 beads. 5% input and the precipitated proteins were analyzed with antibodies against HA and FLAG. (**D**) HeLa cells were transfected with GFP-Stx17 and one of the indicated FLAG-tagged constructs. At 24 hr after transfection, cell lysates were prepared and immunoprecipitated using an anti-GFP antibody. 5% input and the precipitated proteins were analyzed with antibodies against GFP and FLAG. (**E**) HeLa cells were mock-transfected or transfected with siRNA Stx17 (440). After 48 hr, cells were transfected with one of the indicated 3x-HA- or FLAG-tagged constructs, and incubated for 24 hr, and then OA was added. After 16 hr, cells were fixed and stained with Lipid Tox and an antibody against HA or FLAG. The staining intensity of HA or FLAG relative to that of Lipid Tox was plotted. Values are the means ± SEM (n = 3).

**Figure 3-source data 2.** Values, statistics, and exact p values for Figure 3-figure supplement 1E.

**Figure 4-source data 1.** Values, statistics, and exact a p value for Figure 4D.

**Figure 5-source data 1.** Values, statistics, and exact p values for Figure 5B-E.

**Figure 6-source data 1**. Values, statistics, and exact p values for Figure 6B.

**Figure 7-source data 1.** Values, statistics, and exact p values for Figure 7A,B.

## References

Arasaki K, Shimizu H, Mogari H, Nishida N, Hirota N, Furuno A, Kudo Y, Baba M, Baba N, Cheng J, Fujimoto T, Ishihara N, Ortiz-Sandoval C, Barlow LD, Raturi A, Dohmae N, Wakana Y, Inoue H, Tani K, Dacks JB, Simmen T, Tagaya M. 2015. A role for the ancient SNARE syntaxin 17 in regulating mitochondrial division. Developmental Cell 32:304-317. doi: 10.1016/j.devcel.2014.12.011

Boström P, Andersson L, Rutberg M, Perman J, Lidberg U, Johansson BR, Fernandez-Rodriguez J, Ericson J, Nilsson T, Borén J, Olofsson SO. 2007. SNARE proteins mediate fusion between cytosolic lipid droplets and are implicated in insulin sensitivity. Nature Cell Biology 9: 1286-1293.

Brasaemle DL, Dolios G, Shapiro L, Wang R. 2004. Proteomic analysis of proteins associated with lipid droplets of basal and lipolytically stimulated 3T3-L1 adipocytes. The Journal of Biological Chemistry 279: 46835-46842.

Cartwright BR, Binns DD, Hilton CL, Han S, Gao Q, Goodman JM. 2015. Seipin performs dissectible functions in promoting lipid droplet biogenesis and regulating droplet morphology. Molecular Biology of the Cell 26:726-739. doi: 10.1091/mbc.E14-08-1303

Chipuk JE, McStay GP, Bharti A, Kuwana T, Clarke CJ, Siskind LJ, Obeid LM, Green DR. 2012. Sphingolipid metabolism cooperates with BAK and BAX to promote the mitochondrial pathway of apoptosis. Cell 148:988-1000. doi: 10.1016/j.cell.2012.01.038

Choudhary V, Ojha N, Golden A, Prinz WA. 2015. A conserved family of proteins facilitates nascent lipid droplet budding from the ER. The Journal of Cell Biology 211:261-271. doi: 10.1083/jcb.201505067

Diao J, Liu R, Rong Y, Zhao M, Zhang J, Lai Y, Zhou Q, Wilz LM, Li J, Vivona S, Pfuetzner RA, Brunger AT, Zhong Q. 2015. ATG14 promotes membrane tethering and fusion of autophagosomes to endolysosomes. Nature 520:563-566. doi: 10.1038/nature1414

Fei W, Zhong L, Ta MT, Shui G, Wenk MR, Yang H. 2011. The size and phospholipid composition of lipid droplets can influence their proteome. Biochemical and Biophysical Research Communications 415:455-462. doi: 10.1016/j.bbrc.2011.10.091

Fujimoto Y, Itabe H, Kinoshita T, Homma KJ, Onoduka J, Mori M, Yamaguchi S, Makita M, Higashi Y, Yamashita A, Takano T. 2007. Involvement of ACSL in local synthesis of neutral lipids in cytoplasmic lipid droplets in human hepatocyte HuH7. The Journal of Lipid Research 48: 1280-1292.

Fujimoto Y, Itabe H, Sakai J, Makita M, Noda J, Mori M, Higashi Y, Kojima S, Takano T. 2004. Identification of major proteins in the lipid droplet-enriched fraction isolated from the human hepatocyte cell line HuH7. Biochimica et Biophysica Acta 1644: 47-59.

Gao Q, Goodman JM. 2015. Frontier in Cell and Developmental Biology 3:49. doi: 10.3389/fcell.2015.00049

Grippa A, Buxó L, Mora G, Funaya C, Idrissi FZ, Mancuso F, Gomez R, Muntanyà J, Sabidó E, Carvalho P. 2015. The seipin complex Fld1/Ldb16 stabilizes ER-lipid droplet contact sites. The Journal of Cell Biology 211:829-844. doi: 10.1083/jcb.201502070

Hamasaki M, Furuta N, Matsuda A, Nezu A, Yamamoto A, Fujita N, Oomori H, Noda T, Haraguchi T, Hiraoka Y, Amano A, Yoshimori T. 2013. Autophagosomes form at ER-mitochondria contact sites. Nature 495:389-393. doi: 10.1038/nature11910

Hayashi T, Su TP. 2007. Sigma-1 receptor chaperones at the ER-mitochondrion interface regulate Ca2+ signaling and cell survival. Cell 131: 596-610.

Hisanaga Y, Ago H, Nakagawa N, Hamada K, Ida K, Yamamoto M, Hori T, Arii Y, Sugahara M, Kuramitsu S, Yokoyama S, Miyano M. 2004. Structural basis of the substrate specific two-step catalysis of long chain fatty acyl-CoA synthetase dimer. The Journal of Biological Chemistry 279: 31717-31726.

Ingelmo-Torres M, González-Moreno E, Kassan A, Hanzal-Bayer M, Tebar F, Herms A, Grewal T, Hancock JF, Enrich C, Bosch M, Gross SP, Parton RG, Pol A. 2009. Hydrophobic and basic domains target proteins to lipid droplets. Traffic 10:1785-1801. doi:10.1111/j.1600-0854.2009.00994.x

Itakura E, Kishi-Itakura C, Mizushima N. 2012. The hairpin-type tail-anchored SNARE syntaxin 17 targets to autophagosomes for fusion with endosomes/lysosomes. Cell 151:1256-1269. doi: 10.1016/j.cell.2012.11.001

Jägerström S, Polesie S, Wickström Y, Johansson BR, Schröder HD, Højlund K, Boström P. 2009. Lipid droplets interact with mitochondria using SNAP23. Cell Biology International 33:934-940. doi: 10.1016/j.cellbi.2009.06.011

Jacquier N, Choudhary V, Mari M, Toulmay A, Reggiori F, Schneiter R. 2011. Lipid droplets are functionally connected to the endoplasmic reticulum in Saccharomyces cerevisiae. Journal of Cell Science 124:2424-2437. doi: 10.1242/jcs.076836

Kassan A, Herms A, Fernández-Vidal A, Bosch M, Schieber NL, Reddy BJ, Fajardo A, Gelabert-Baldrich M, Tebar F, Enrich C, Gross SP, Parton RG, Pol A. 2013. Acyl-CoA synthetase 3 promotes lipid droplet biogenesis in ER microdomains. The Journal of Cell Biology 203:985-1001. doi: 10.1083/jcb.201305142

Kory N, Farese RV Jr, Walther TC. 2016. Targeting Fat: Mechanisms of Protein Localization to Lipid Droplets. Trends in Cell Biology 26:535-546. doi: 10.1016/j.tcb.2016.02.007

Kuerschner L, Moessinger C, Thiele C. 2008. Imaging of lipid biosynthesis: how a neutral lipid enters lipid droplets. Traffic 9: 338-352.

Longo PA, Kavran JM, Kim MS, Leahy DJ. (2013). Transient mammalian cell transfection with polyethylenimine (PEI). Methods in Enzymology 529:227-240. doi: 10.1016/B978-0-12-418687-3.00018-5

Lynes EM, Bui M, Yap MC, Benson MD, Schneider B, Ellgaard L, Berthiaume LG, Simmen T. 2012, Palmitoylated TMX and calnexin target to the mitochondria-associated membrane. The EMBO Journal 31:457-470. doi: 10.1038/emboj.2011.384

Magré J, Delépine M, Khallouf E, Gedde-Dahl T Jr, Van Maldergem L, Sobel E, Papp J, Meier M, Mégarbané A, Bachy A, Verloes A, d'Abronzo FH, Seemanova E, Assan R, Baudic N, Bourut C, Czernichow P, Huet F, Grigorescu F, de Kerdanet M, et al. 2001. Identification of the gene altered in Berardinelli-Seip congenital lipodystrophy on chromosome 11q13. Nature Genetics 28: 365-370.

McLelland GL, Lee SA, McBride HM, Fon EA. 2016. Syntaxin-17 delivers PINK1/parkin-dependent mitochondrial vesicles to the endolysosomal system. The Journal of Cell Biology 214:275-91. doi: 10.1083/jcb.201603105

Moessinger C, Klizaite K, Steinhagen A, Philippou-Massier J, Shevchenko A, Hoch M, Ejsing CS, Thiele C. 2014. Two different pathways of phosphatidylcholine synthesis, the Kennedy Pathway and the Lands Cycle, differentially regulate cellular triacylglycerol storage. BMC Cell Biology 15:43. doi: 10.1186/s12860-014-0043-3

Moessinger C, Kuerschner L, Spandl J, Shevchenko A, Thiele C. 2011. Human lysophosphatidylcholine acyltransferases 1 and 2 are located in lipid droplets where they catalyze the formation of phosphatidylcholine. The Journal of Biological Chemistry 286:21330-21339. doi: 10.1074/jbc.M110.202424

Myhill, N., Lynes, E.M., Nanji, J.A., Blagoveshchenskaya, A.D., Fei, H., Carmine Simmen, K., Cooper, T.J., Thomas, G., and Simmen, T. (2008). The subcellular distribution of calnexin is mediated by PACS-2. Molecular Biology of the Cell 19:2777-2788. doi: 10.1091/mbc.E07-10-0995

Naon D, Zaninello M, Giacomello M, Varanita T, Grespi F, Lakshminaranayan S, Serafini A, Semenzato M, Herkenne S, Hernández-Alvarez MI, Zorzano A, De Stefani D, Dorn GW 2nd, Scorrano L. 2016. Critical reappraisal confirms that Mitofusin 2 is an endoplasmic reticulum-mitochondria tether. Proceedings of the National Academy of Sciences of the United States of America 113: 11249-11254.

Ohsaki Y, Kawai T, Yoshikawa Y, Cheng J, Jokitalo E, Fujimoto T. 2016. PML isoform II plays a critical role in nuclear lipid droplet formation. The Journal of Cell Biology 212:29-38. doi: 10.1083/jcb.201507122

Ohsaki Y, Suzuki M, Fujimoto T. 2017. The lipid droplet and the endoplasmic reticulum. In Organelle contact sites: from molecular mechanism to disease. Tagaya M and Simmen T (Editors) In Advances in Experimental Medicine and Biology vol. 997 Switzerland: Springer, in press.

Ohsaki Y, Suzuki M, Fujimoto T. 2014. Open questions in lipid droplet biology. Chemistry and Biology 21:86-96. doi: 10.1016/j.chembiol.2013.08.009

Oliferenko S, Paiha K, Harder T, Gerke V, Schwärzler C, Schwarz H, Beug H, Günthert U, Huber LA. 1999. Analysis of CD44-containing lipid rafts: Recruitment of annexin II and stabilization by the actin cytoskeleton. The Journal of Cell Biology 146: 843-854.

Pagac M, Cooper DE, Qi Y, Lukmantara IE, Mak HY, Wu Z, Tian Y, Liu Z, Lei M, Du X, Ferguson C, Kotevski D, Sadowski P, Chen W, Boroda S, Harris TE, Liu G, Parton RG, Huang X, Coleman RA, Yang H. 2016. SEIPIN regulates lipid droplet expansion and adipocyte development by modulating the activity of glycerol-3-phosphate acyltransferase. Cell Reports 217:1546-1559. doi: 10.1016/j.celrep.2016.10.037

Pol A, Gross SP, Parton RG. 2014. Biogenesis of the multifunctional lipid droplet: lipids, proteins, and sites. The Journal of Cell Biology 204:635-646. doi: 10.1083/jcb.201311051

Poppelreuther M, Rudolph B, Du C, Großmann R, Becker M, Thiele C, Ehehalt R, Füllekrug J. 2012. The N-terminal region of acyl-CoA synthetase 3 is essential for both the localization on lipid droplets and the function in fatty acid uptake. The Journal of Lipid Research 53:888-900. doi: 10.1194/jlr.M024562

Rambold AS, Cohen S, Lippincott-Schwartz J. 2015. Fatty acid trafficking in starved cells: regulation by lipid droplet lipolysis, autophagy, and mitochondrial fusion dynamics. Developmental Cell 32:678-692. doi: 10.1016/j.devcel.2015.01.029

Rusiñol AE, Cui Z, Chen MH, Vance JE. 1994. A unique mitochondria-associated membrane fraction from rat liver has a high capacity for lipid synthesis and contains pre-Golgi secretory proteins including nascent lipoproteins. The Journal of Biological Chemistry 269: 27494-27502.

Salaün C, Gould GW, Chamberlain LH. 2005. The SNARE proteins SNAP-25 and SNAP-23 display different affinities for lipid rafts in PC12 cells. Regulation by distinct cysteine-rich domains. The Journal of Biological Chemistry 280: 1236-1240.

Salaun C, Greaves J, Chamberlain LH. 2010. The intracellular dynamic of protein palmitoylation. The Journal of Cell Biology 91:1229-1238. doi: 10.1083/jcb.201008160.

Salo VT, Belevich I, Li S, Karhinen L, Vihinen H, Vigouroux C, Magré J, Thiele C, Hölttä-Vuori M, Jokitalo E, Ikonen E. 2016. Seipin regulates ER-lipid droplet contacts and cargo delivery. The EMBO Journal 35: 2699-2716.

Sano R, Annunziata I, Patterson A, Moshiach S, Gomero E, Opferman J, Forte M, d’Azzo A. 2009. GM1-ganglioside accumulation at the mitochondria-associated ER membranes links ER stress to Ca2+-dependent mitochondrial apoptosis. Molecular Cell 36:500-511. doi: 10.1016/j.molcel.2009.10.021

Shindou H, Shimizu T. 2009. Acyl-CoA:lysophospholipid acyltransferases. The Journal of Biological Chemistry 284:1-5. doi: 10.1074/jbc.R800046200

Simmen T, Aslan JE, Blagoveshchenskaya AD, Thomas L, Wan L, Xiang Y, Feliciangeli SF, Hung CH, Crump CM, Thomas G. 2005. PACS-2 controls endoplasmic reticulum-mitochondria communication and Bid-mediated apoptosis. The EMBO Journal 24: 717-729.

Somwar R, Roberts CT Jr, Varlamov O. 2011. Live-cell imaging demonstrates rapid cargo exchange between lipid droplets in adipocytes. FEBS Letters 585:1946-1950. doi: 10.1016/j.febslet.2011.05.016.

Soupene E, Dinh NP, Siliakus M, Kuypers FA. 2010. Activity of the acyl-CoA synthetase ACSL6 isoforms: role of the fatty acid Gate-domains. <u>BMC Biochemstry</u> 11:18. doi: 10.1186/1471-2091-11-18

Steegmaier M, Oorschot V, Klumperman J, Scheller RH. 2000. Syntaxin 17 is abundant in steroidogenic cells and implicated in smooth endoplasmic reticulum membrane dynamics. Molecular Biology of the Cell 11: 2719-2731.

Stone SJ, Levin MC, Zhou P, Han J, Walther TC, Farese RV Jr. 2009. The endoplasmic reticulum enzyme DGAT2 is found in mitochondria-associated membranes and has a mitochondrial targeting signal that promotes its association with mitochondria. The Journal of Biological Chemistry 284:5352-5361. doi: 10.1074/jbc.M80576820

Stordeur C, Puth K, Sáenz JP, Ernst R. 2014. Crosstalk of lipid and protein homeostasis to maintain membrane function. Biological Chemistry 395:313-326. doi: 10.1515/hsz-2013-0235

Szymanski KM, Binns D, Bartz R, Grishin NV, Li WP, Agarwal AK, Garg A, Anderson RG, Goodman JM. 2007. Proceedings of the National Academy of Sciences of the United States of America 104: 20890-20895.

Takáts S, Nagy P, Varga Á, Pircs K, Kárpáti M, Varga K, Kovács AL, Hegedűs K, Juhász G. 2013. Autophagosomal Syntaxin17-dependent lysosomal degradation maintains neuronal function in Drosophila. The Journal of Cell Biology 201:531-539. doi: 10.1083/jcb.201211160

Thiam AR, Beller M. 2017. The why, when and how of lipid droplet diversity. Journal of Cell Science 130:315-324. doi: 10.1242/jcs.192021

Thiam AR, Forêt L. 2016. The physics of lipid droplet nucleation, growth and budding. Biochimica et Biophysica Acta 1861:715-722. doi: 10.1016/j.bbalip.2016.04.018

Vance JE. 2014, MAM (mitochondria-associated membranes) in mammalian cells: lipids and beyond. Biochimica et Biophysica Acta 1841:595-609. doi: 10.1016/j.bbalip.2013.11.014

Vogel K, Roche PA. 1999. SNAP-23 and SNAP-25 are palmitoylated in vivo. Biochemical and Biophysical Research Communications. 258: 407-410.

Walther TC, Farese RV Jr. 2012. Lipid droplets and cellular lipid metabolism. Annual Review of Biochemistry 81:687-714. doi: 10.1146/annurev-biochem-061009-102430

Wang CW, Miao YH, Chang YS. 2014. Control of lipid droplet size in budding yeast requires the collaboration between Fld1 and Ldb16. Journal of Cell Science 127:1214-28. doi: 10.1242/jcs.1377

Wang H, Becuwe M, Housden BE, Chitraju C, Porras AJ, Graham MM, Liu XN, Thiam AR, Savage DB, Agarwal AK, Garg A, Olarte MJ, Lin Q, Fröhlich F, Hannibal-Bach HK, Upadhyayula S, Perrimon N, Kirchhausen T, Ejsing CS, Walther TC, Farese RV. 2016. Seipin is required for converting nascent to mature lipid droplets. Elife 5. pii: e16582. doi: 10.7554/eLife.16582

Welte MA. 2015. Expanding roles for lipid droplets. Current Biology 25:R470-481. doi:10.1016/j.cub.2015.04.00

Wilfling F, Wang H, Haas JT, Krahmer N, Gould TJ, Uchida A, Cheng JX, Graham M, Christiano R, Fröhlich F, Liu X, Buhman KK, Coleman RA, Bewersdorf J, Farese RV Jr, Walther TC. 2013. Triacylglycerol synthesis enzymes mediate lipid droplet growth by relocalizing from the ER to lipid droplets. Developmental Cell 24:384-399. doi: 10.1016/j.devcel.2013.01.013

Zehmer JK., Bartz R, Liu P, Anderson RG. 2008. Identification of a novel N-terminal hydrophobic sequence that targets proteins to lipid droplets. Journal of Cell Science 121:1852-1860. doi: 10.1242/jcs.012013

